# IOBR2: Multidimensional Decoding Tumor Microenvironment for Immuno-Oncology Research

**DOI:** 10.1101/2024.01.13.575484

**Authors:** Dongqiang Zeng, Yiran Fang, Peng Luo, Wenjun Qiu, Shixiang Wang, Rongfang Shen, Wenchao Gu, Xiatong Huang, Qianqian Mao, Yonghong Lai, Xi Xu, Min Shi, Guangchuang Yu, Wangjun Liao

## Abstract

The use of large transcriptome datasets has greatly improved our understanding of the tumor microenvironment (TME) and helped develop precise immunotherapies. The increasing popularity of multi-omics sequencing, single-cell transcriptome sequencing (scRNA), and spatial transcriptome sequencing has led to numerous new discoveries. However, these findings require clinical phenotypic validation with a large sample size. To enhance the integration of multi-omics in advancing research on the tumor microenvironment, we have developed a systematic and comprehensive analytical tool (Immuno-Oncology Biological Research 2, IOBR2) based on our prior work. IOBR2 offers six modules for TME analysis based on multi-omics data. These modules cover data preprocessing, TME estimation, TME infiltrating patterns, cellular interactions, genome and TME interaction, and visualization for TME relevant features, as well as modelling based on key features. IOBR2 integrates multiple vital microenvironmental analysis algorithms and signature estimation methods, simplifying the analysis and downstream visualization of the TME. In addition to providing a quick and easy way to construct gene signatures from single-cell data, IOBR2 also provides a way to construct a reference matrix for TME deconvolution from single-cell RNAseq. The analysis pipeline and feature visualization are user-friendly and provide a comprehensive description of the complex TME, offering insights into tumor-immune interactions. A comprehensive gitbook (https://iobr.github.io/book/) is available with a user-friendly manual and complete analysis workflow for each module.

## Background

Studies of the tumour microenvironment using gene expression patterns from large bulk transcriptome datasets have advanced the understanding and identification of the interactions within microenvironment and aided in the development of more precise immunotherapy treatments for tumours. This progress includes the utilization of gene signature scores, such as T cell-inflamed gene expression profile (GEP) [1], pan-fibroblast TGFβ response signature (Pan-F-TBRS) [2], tertiary lymphoid structures (TLS) [3], and TMEscore [4], as well as MFP, a model based on machine learning. Years of research have shown that transcriptome gene expression signatures can adeptly characterise the tumour microenvironment and exhibit significant clinical translational potential [5]. IOBR (Immune-Oncology Biological Research) was debuted in 2021 to describe the systematic approach to tumour microenvironment profile and correlation [6]. This tool has enabled numerous studies to come to fruition over the last few years. Simultaneously, we are continuously enhancing and updating IOBR with the assistance of our users. The recent surge in single-cell RNA sequencing (scRNA-seq) has enabled us to identify novel microenvironmental cells, tumour microenvironmental characteristics, and tumour clonal signatures with higher accuracy [7]. It is necessary to scrutinize, confirm and depict these features attained from high-dimensional single-cell information in bulk-seq with extended specimen sizes for clinical phenotyping. We have developed a systematic analysis process for IOBR based on the needs of the aforementioned studies.

Our team has developed a suit of highly effective functions for microenvironmental analysis. Furthermore, we have integrated 8 key microenvironmental analysis algorithms into our framework, including CIBERSORT [8], EPIC [9] and quanTIseq [10]. This integration makes it simple for users to conduct analyses and visualize data using the IOBR process. However, the identification of new cells and functions has posed several challenges to users attempting to customize parsing with newly acquired reference data. The advancement in artificial intelligence and machine learning are driving researchers to focus on identifying patterns within tumour microenvironments and exploring the clinical importance of microenvironmental features [11]. In addition, screening important features, evaluating feature robustness, and constructing models have become pressing concerns. To tackle these challenges, we compile additional algorithms with the aim of creating a multidimensional analysis and visualization procedure focused on parsing data concerning the tumour microenvironment.

Through the establishment of systematic modules for tumour microenvironment analysis, we have successfully conducted multi-dimensional analyses of the tumour microenvironment. This involves data quality control and processing, parsing the tumour microenvironment, exploring interactions within the tumour microenvironment, as well as the interactions between the microenvironment and its genome, visualizing the tumour microenvironment, and features screening and constructing models based on key features. We have created a comprehensive gitbook with a user-friendly manual for researchers (https://iobr.github.io/book/). This system is perfect for large-scale research in multi-omics related to the tumour microenvironment. It offers a valuable and reproducible method for enhancing precision in diagnosis and treatment, guided by the tumour microenvironment. Additionally, it enables experts to discover new therapeutic targets and overcome tumour therapeutic resistance.

## Method

### The framework of IOBR

Building upon the existing functions of IOBR, IOBR2 introduces additional analysis and visualization capabilities, with its comprehensive implementation and functionalities thoroughly detailed in the tutorial (https://iobr.github.io/book/) with a complete analysis pipeline [6]. The current version, IOBR 2.0, encompasses six functional modules: 1) Transcriptome data prepare module (pre-procession of transcriptome data, as well as pertinent batch statistical analyses); 2) TME deconvolution and signature estimation module (estimation of signature scores and identification of phenotype relevant signatures, along with decoding immune contexture); 3) TME interaction module (clustering TME characteristics and analyzing receptor-ligand interactions); 4) Genome and TME interaction module (analysis of signature associated mutations); 5) TME data visualization and Statistical analysis module (visual representation and statistical examination of TME data); 6) TME modeling module (fast model construction and the assessment of model performance).

### Transcriptome data prepare

#### Data preparation

In line with the preprocessing workflow of transcriptomic data, we have integrated a variety of functionalities into the IOBR2. IOBR2 supports users in retaining genes based on the maximum or average values of repeated gene expressions. Additionally, we have developed an annotation function for annotating expression matrices. The annotation files in IOBR include *anno_hug133plus2, anno_rnaseq*, and *anno_illumina*, corresponding to annotations for HG-U133 Plus 2.0 microarray probes, RNAseq annotation data, and Illumina microarray probes, respectively.

In IOBR2, we have established a function for differentially expressed genes (DEGs) analysis between two groups. This function supports two analytical methods, limma [12] and Desq2 [13]. The limma employs a linear model to assess changes in gene expression, correcting for multiple testing differences using an empirical Bayesian method. Originally designed for microarray data, its utility has been extended to small-sample RNA-seq data analysis. Desq2, specifically designed for RNA-seq data analysis, uses a negative binomial distribution to model gene expression data, applying either the Wald test or likelihood ratio test to each gene to detect expression differences. Users can choose the appropriate method based on their data type and research needs. Additionally, IOBR2 supports DEG analysis for more than 2 groups. It leverages the Seurat R package to identify significant markers across multiple groups within the dataset [14]. The methods available for comparison include bootstrap, delong, and venkatraman, offering a range of options for comprehensive analysis.

Users can also rapidly convert gene expression count data into Transcripts Per Million (TPM) value. During the annotation and conversion processes, additional operations such as merging annotation data with the expression matrices, removing unnecessary columns, transforming rows and columns, and handling duplicates based on the specified method can be simultaneously implemented. For sequencing data from different sources or batches, users can use IOBR to examine batch effects in the data and perform batch correction. Furthermore, we have built a filtering function rapid analysis of gene expression data and identification of outliers in the dataset.

### TME deconvolution and signature estimation

#### Signature Estimation

To enhance the characterization of the TME in cancer cells and to deepen our understanding of tumor immunity and its functional states, we have developed an estimation function for user-generated signatures or 323 reported signatures enrolled in IOBR (**Supplementary Table S1**). The extensive signature collection is categorized into three distinct groups: TME-associated, tumor-metabolism, and tumor-intrinsic signatures. Additionally, IOBR supports the estimation of the signature gene sets derived from the GO, KEGG, HALLMARK, and REACTOME databases. IOBR allows users to generate custom signature lists aligned with their own biological discovery or exploratory needs, thereby streamlining the estimation process and enabling systematic follow-up exploration. Users also have the option to generate signature lists from single-cell differential analysis or the Msigdb database (gsea-msigdb.org) for their subsequent research.

In the evaluation of signature scores, we incorporated three methodologies: Single-sample Gene Set Enrichment Analysis (ssGSEA), Principal Component Analysis (PCA), and Z-score. ssGSEA is extensively used to evaluate the enrichment or activity of specific gene sets within individual samples [15]. Each ssGSEA enrichment score reflects the collective expression dynamics of a specific gene set in a single sample, indicating whether the genes in the set are collectively upregulated or downregulated in expression. Significantly, PCA calculates the principal components to reduce the dimensionality of data simultaneously preserving the maximum variability of data for predictive model construction. Current signatures constructed using PCA methodology include the Pan-F-TBRs [2] and the TMEscore [4, 16], two promising biomarkers for predicting clinical outcomes and assessing the sensitivity of malignancies to treatments. Z-score, a statistical metric, measures a score’s deviation from the mean of a dataset in standard deviations.

#### TME Deconvolution

Different mechanisms in the tumor microenvironment (TME) are involved in mediating the immune response and affect the efficacy of treatment. The important aspect is the cell-type composition of the TME, which is the key elements shaping the intricate landscape of anti-tumor immunity [5]. Deciphering the cellular composition of the TME is a significant technical challenge, addressed by various deconvolution algorithms, each with its unique advantages and limitations [17, 18]. IOBR integrates eight open-source deconvolution methodologies: CIBERSORT [8], ESTIMATE [19], quanTIseq [10], TIMER [20], IPS [21], MCPCounter [22], xCell [23], and EPIC [9].

CIBERSORT is the most well-recognized method for identifies 22 immune cell types in TME, allowing large-scale analysis of RNA mixtures for cellular biomarkers and therapeutic targets with promising accuracy [8]. Notably, IOBR leverages CIBERSORT’s linear vector regression principle, allowing users to create custom signatures and extending its input file compatibility to cell subsets derived from single-cell sequencing results. ESTIMATE focuses on non-malignant components, like stromal and immune signatures, to assess tumor purity [19]. The quanTIseq method quantifies 10 immune cell subsets from bulk RNA-seq data [10]. TIMER is adept at quantifying the abundance of tumor-infiltrating immune compartments [20]. It offers six major analytic modules, enabling detailed analysis of immune infiltration alongside other cancer molecular profiles. IPS calculates 28 TIL subpopulations, including effector and memory T cells and immunosuppressive cells [21]. MCP-counter robustly quantifies the absolute abundance of eight immune and two stromal cell populations within heterogeneous tissues, using transcriptomic data [22]. xCell offers an extensive analysis of 64 immune cell types from RNA-seq data, including various cell subsets in bulk tumor tissues [23]. EPIC decodes the proportion of immune and cancer cells from the expression of genes, comparing it to specific cell expression profiles to accurately predict the cellular subpopulation landscape [9].

#### Signatures Derived From scRNA-seq Data

Moving beyond traditional bulk sequencing, validating the clinical relevance of cell types identified by scRNA-seq becomes essential. To facilitate this, IOBR integrates CIBERSORT’s linear Support Vector Regression (SVR) with the Least Squares Estimate of Imbalance (LSEI) algorithms [24], enabling a streamlined analysis of bulk RNA-seq data for the clinical validation of targets identified through scRNA-seq data. In addition, IOBR2 allows the user to screen DEGs of cell-types based on “sce” projects generated by the Seurat package [14], performing cellular deconvolution related to the research objectives or constructing signature gene lists for IOBR2 scoring calculations.

#### TME interaction module

To facilitate a deeper analysis of the TME and identify distinct TME patterns in patients, we have developed a clustering function based on the NbClust R package. This function enables unsupervised clustering analysis using datasets generated by users or signature estimation scores from IOBR. Based on the results, IOBR can determine the optimal number of clusters and assign each sample to a specific cluster.

Additionally, IOBR offers a function for analyzing ligand-receptor pairs within the TME. It evaluates 813 pairs of ligand-receptor interactions based on gene expression patterns. These pairs are expressed in 25 cell types that are present in the TME, encompassing immune cells, cancer cells, fibroblasts, endothelial cells, and adipocytes [25]. Users provide transcriptomic data as input, allowing IOBR to generate group-specific, system-based signatures of the TME. A pairwise Wilcoxon test is then employed to identify distinctive signatures and ligand-receptor interactions between different groups.

#### Genome and TME interaction

IOBR not only focuses on systematic signature-phenotype studies but also expands its research scope to include the exploration of interactions between transcriptomes, microenvironments, and genome profiles. It accepts genome data in Mutation Annotation Format (MAF) [26] downloaded from the UCSC website or user-generated mutation matrices as input for identifying mutations associated with specific signatures. Additionally, IOBR supports the transformation of MAF data into a comprehensive mutation matrix. This matrix contains data on distinct variation types, including insertion-deletion mutations (indels), single-nucleotide polymorphisms (SNPs), and frameshift mutations, or it can integrate all these mutation types, offering users flexible selection options. For the analysis of mutations significantly linked to targeted signatures, IOBR employs the Wilcoxon rank-sum test in this module for batch analysis. Moreover, IOBR supports batch visualization, allowing users to easily view and interpret the mutation status (mutation or non-mutation) of specified genes or regions.

#### TME data visualization and Statistical analysis

Batch analysis and visualization of results from the TME deconvolution and signature estimation module are pivotal features of IOBR2. To implement TME deconvolution and signature computation for potential clinical translation, we have systematically categorized the collected signatures into 43 groups (**Supplementary Table S2**), expanding upon the foundation of IOBR. These categories encompass TME cell populations (classified by deconvolution methods, cell types, or scRNA-seq results), signatures of immune phenotypes, tumor metabolism, HALLMARK and so on. Users can freely adjust the number of signatures within each group and also utilize signatures documented in IOBR2 to construct new groups for research exploration. IOBR2 also supports the construction of new signature groups based on immune-oncological research findings or specific study objectives, enabling users to tailor their analysis to their unique research needs. Further, we integrate a visualization function specifically for batch correlation analysis of signature groups, either user-generated or enrolled in IOBR2. This function allows for visualizations based on specified groups, including boxplots and heatmaps, and employs the Wilcoxon Rank Sum Test to compare statistical differences in signatures between groups. Moreover, IOBR2 is capable of presenting TME cell fractions as percentage bar charts in batch visualization, supporting input of deconvolution results from “CIBERSORT”, “EPIC” and “quanTIseq” methodologies to further compare the TME cell distributions within one sample or among different samples.

To provide a more intuitive understanding of the TME and streamline the analysis process, IOBR2 has introduced a range of new batch visualization and statistical functions. The batch analysis methods supported by IOBR2 include batch Wilcoxon rank-sum test between two groups,batch calculation of hazard ratios and confidence intervals for the specified signature, and batch analysis of correlation. IOBR2 supports computing correlations between two features or genes, or between a target variable and multiple variables, visualizing these correlations through heatmaps or scatter plots. It supports two correlation methods: Pearson correlation coefficient or Spearman’s rank correlation coefficient Furthermore, IOBR2 provides other independent analysis and visualization functions, including KM survival analysis, GSEA, and PCA analysis. Notably, IOBR2 allows users to perform GSEA based on user-generated signatures or signatures registered in IOBR2.

The TME data visualization and statistical analysis module of IOBR2 collectively enable easy integration and visualization of the aforementioned deconvolution results, offering flexibility in selecting specific methodologies of interest. This module permits systematic identification of phenotype-relevant signatures, cell fractions, or signature genes, accompanied by corresponding batch statistical analyses and visualization options. Within IOBR2, these methods are available for users to choose for targeted analysis or integration, complemented by a range of visualization tools.

#### TME modeling

For effective application of the signatures in clinical interpretation, IOBR2 provides functions for feature selection, robust biomarker identification, and model construction based on prior identified phenotype associated signatures. Utilizing these features to build prognostic models holds promise for accurately and cost-effectively predicting tumor patients’ survival and treatment sensitivities. In addition, IOBR2 supports the performance assessment of models predicting patient survival and treatment responses, offering valuable tools for evaluating the efficacy and applicability of these models in clinical settings.

#### Availability of data and materials

IOBR R package can be available at https://github.com/IOBR/IOBR.

## Results

To fully utilize transcriptomic data in uncovering immune-tumor interactions and their potential clinical applications, we have expanded the capabilities of IOBR. Beyond integrating conventional analysis and deep mining methods, we have extended our functionalities to include TME interaction analysis, providing a comprehensive one-stop solution for analysis and visualization in transcriptome projects.

### IOBR workflow

Based on prior research, we have enhanced the analytical and visualization capabilities in IOBR2. Utilizing these advancements, we have structured the analytical workflow of IOBR into six functional modules: Transcriptome data preparation module, TME deconvolution and signature estimation module, TME interaction module, Genome and TME interaction module, TME data visualization and Statistical analysis module, and TME modeling module. The schematic workflow and functional codes are depicted in Figures 1, 2, respectively. Corresponding figures were dynamically generated following inputting function-specific parameters of pertinent modules. Details of these six modules are illustrated in the Methods sections. Charts derived from IOBR reach quality requirements of publication and can be flexibly modified locally. Drawing from previously published studies, we have incorporated several example datasets into IOBR. The workflow and functionalities of IOBR are further illustrated below using real-world data from these datasets.

**Fig. 1.**
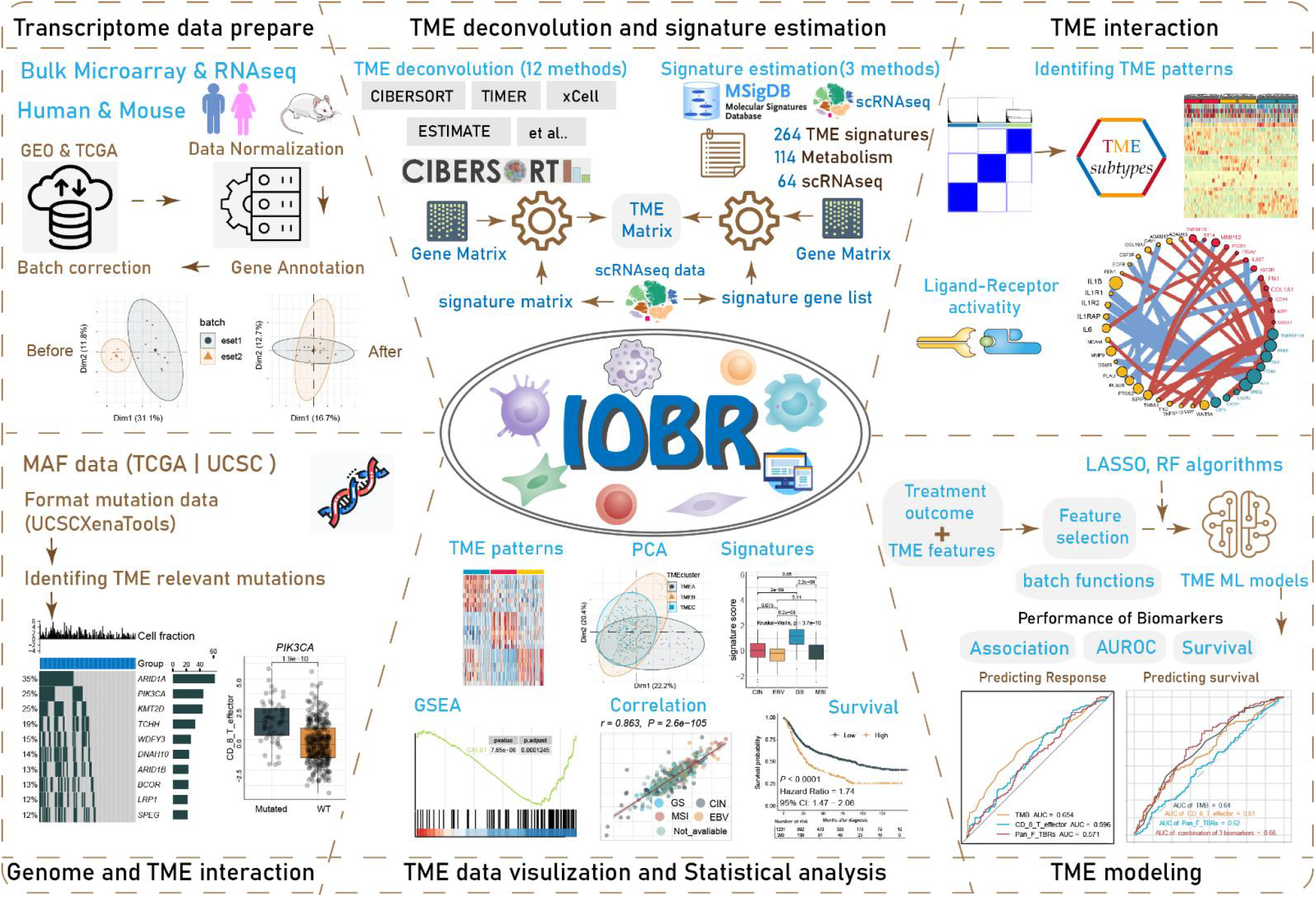
The graphical scheme describing the workflow of the IOBR2. IOBR2 encompasses transcriptome data preparation, multiple deconvolution algorithms and signature estimation methods for microenvironment analysis, TME pattern identification, analysis of interactions between the genome and TME, batch visualization and statistical analysis, as well as TME modeling.

**Fig. 2.**
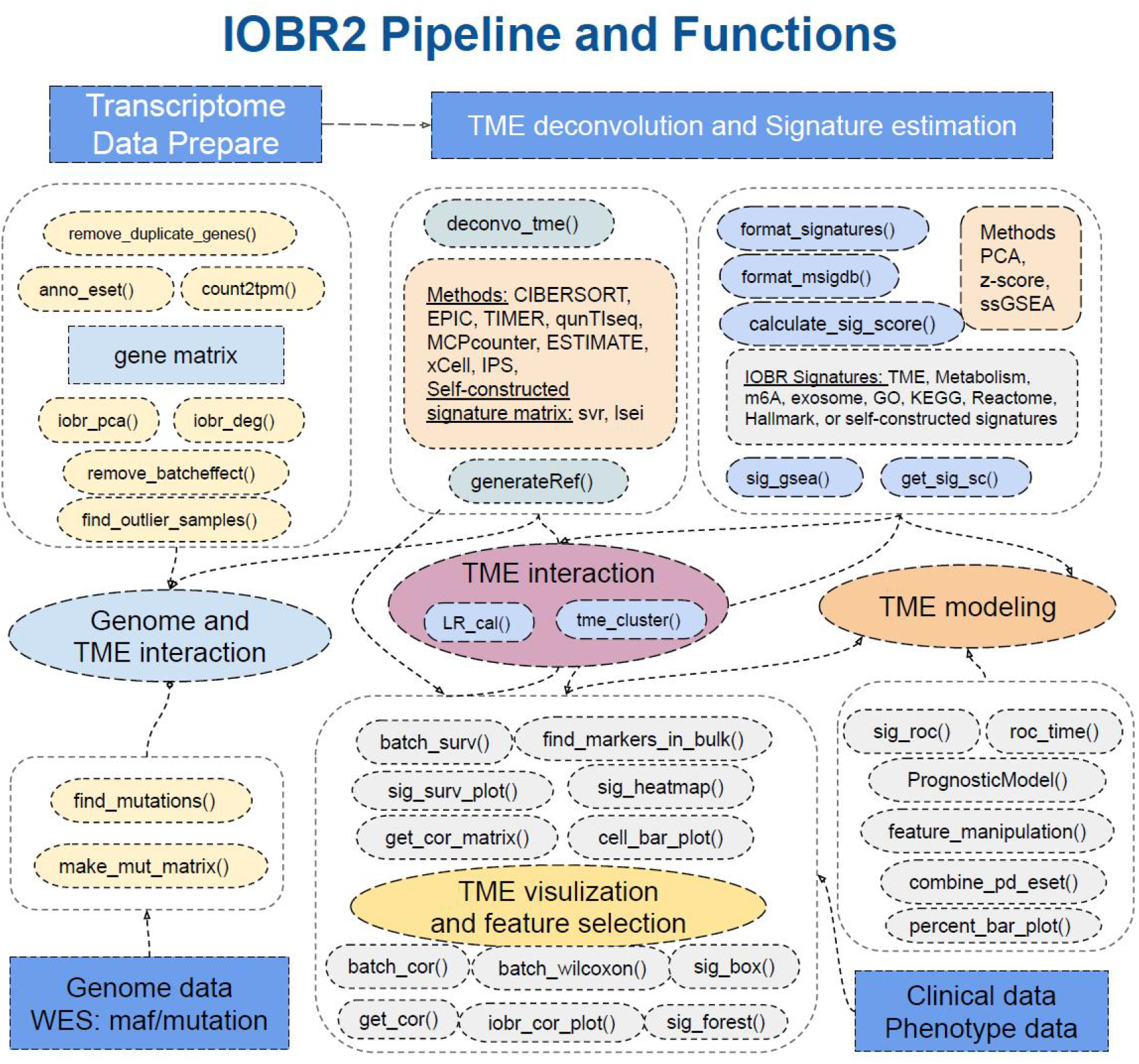
IOBR2 is comprised of six analytic modules related to data preprocess and tumor immune microenvironment. The functionalities of these modules include (1) preprocessing of transcriptome data; (2) estimation of signature scores and identification of phenotype-relevant or user-constructed signatures, along with decoding of TME contexture; (3) identification of TME patterns and analysis of ligand-receptor interactions; (4) estimation of the specific mutation landscape associated with the signature of interest; (5) corresponding batch visualization and statistical analyses; and (6) model construction.

### Multi-group Transcriptome data prepare and DEG analysis

In multi-group transcriptomic studies, preprocessing of sequencing data remains a pivotal focus at the outset of our analysis. For transcriptome sequencing data originating from various sequencing technologies or batches, it’s essential to preprocess the data following RNA-seq read alignment and gene quantification, ensuring comparability across different datasets.

The *remove_duplicate_genes* and *anno_eset* functions eliminate redundant genes and annotate them, allowing for the retention of duplicates based on either average or maximum values, depending on the data characteristics. To mitigate the impact of outliers on subsequent analyses, the *find_outlier_sample* function is used to remove outliers from the annotated data [27].

For transcriptome data derived from different sequencing technologies, batch effects can influence downstream cohort-level analyses. The iobr_pca function, developed on the FactoMineR R package, facilitates principal component analysis (PCA) of all samples, visualizing the results to identify variation patterns across different groups or batches. The remove_batcheffect function, built on the ComBat function from the sva R package, is designed to eliminate batch effects across datasets when analyzing the tumor microenvironment [28].

Differential expression analysis is crucial for identifying genes and pathways that behave differently under specific biological conditions. Following gene annotation and batch effect removal, *iobr_deg* enables differential gene expression analysis between two sample groups. This function employs methods including *Desq2* and *limma* [12, 13] – *Desq2* for RNA-seq data and *limma* for microarray data – and visually presents differences between groups through volcano plots or heatmaps, aiding in the identification of characteristic genes in the data. Further, RNA-seq count data are converted into TPM using the count2tpm function for subsequent analyses.

In summary, IOBR significantly streamlines the data preprocessing process and offers tailored solutions for transcriptomic data from different batches or sequencing technologies. The *iobr_pca* and *remove_batcheffect* functions facilitate visualization of sample clustering and batch effect removal. *iobr_deg*, utilizing *Desq2* and *limma*, conducts differential gene expression analysis between samples.

### Elucidating TME Compositions and Gene Signatures Relevant to Therapeutic Outcomes

Contrasting with many existing approaches that calculate a single signature through a defined method post-analysis, IOBR stands out in its ability to concurrently discern a series of gene signatures, either published or user-generated, utilizing a variety of methodological choices. This capability is instrumental in effectively mapping the interactions between the immune system and tumors. In this context, we extend the application of IOBR to unravel the TME landscape within the growing body of bulk RNA-seq data.

Users can analyze different cell fractions within the Tumor Microenvironment (TME) and estimate multiple gene signatures collected in IOBR using the *deconvo_tme* and *calculate_sig_score* functions.The deconvo_tme function incorporates eight cell fraction deconvolution methods (as detailed in our methodology), aiding in the identification and quantification of various cell types’ relative abundance in the TME. This process reveals the distribution and status of immune cells within the tumor milieu. Additionally, IOBR has amassed a diverse range of gene signatures, including those related to the TME, metabolism, m6A, and others (see methodology), while the *calculate_sig_score* function integrates ssGSEA, PCA, and z-score methods for gene signature scoring in different contexts, allowing researchers to assess patients’ gene expression characteristics from various perspectives. Moreover, users can create customized gene signature lists for *calculate_sig_score* using *format_signatures* and *format_msigdb*, enabling more flexible analysis of transcriptomic data based on their research objectives. Further enhancing its capabilities, IOBR includes the *iobr_cor_plot* function, which visualizes the relationship between different phenotypes and cell types in the TME and their response to treatment using box plots and heatmaps.

Consequently, IOBR amalgamates a variety of deconvolution and scoring algorithms, streamlining the analysis of different data phenotypes and cell type percentages. This facilitates a deeper understanding of gene expression patterns, cellular distributions, and their roles in the context of tumor therapy.

### Leveraging Signature Generated From scRNA-seq Analyses to Unravel bulk-seq data

Current single-cell analyses have unveiled substantial intratumoral cell state heterogeneity at an exceptionally detailed level. Despite this, due to its straightforward methodology and cost-effectiveness, bulk RNA-seq data continue to dominate as the primary approach for gene expression analysis and gene signature assessment. The IOBR tool enables users to apply insights from single-cell derived signatures to examine tumor heterogeneity through bulk RNA-seq data.

Utilizing scRNA-seq data, IOBR allows users to identify cell-type-specific gene expression signatures from cluster analyses reported in existing studies. Marker genes for each cell type are accurately identified through differential expression analysis using the *GenerateRef* function within IOBR. Then, based on the prior knowledge of cell-type-specific gene expression signatures, we can interpret bulk RNA-seq data by utilizing the *deconvo_tme* function, which implements either the linear *svr* algorithm from CIBERSORT or the *lsei* algorithm. Additionally, we employ the *get_sig_sc* function to acquire marker gene objects from single-cell differential analysis, serving as inputs for the *calculate_sig_score* function, thereby enabling the calculation of corresponding signature scores. IOBR is adept at enumerating TME populations, thereby validating findings from single-cell studies and unveiling potential new clinical insights in the context of bulk RNA-seq data. This application effectively links the expanding field of single-cell sequencing research with practical biological insights.

### Identifying TME patterns and analyzing intercellular interactions

The efficacy of a patient’s treatment response is influenced not only by the infiltration of different cell types in the microenvironment but also by the intricate network of interactions among these cells. Hence, to thoroughly dissect the mechanisms underlying patient treatment responses and ultimately predict treatment outcomes, IOBR enables patient classification based on cell infiltration and gene signatures. It also facilitates the analysis of cellular interactions at the bulk RNA-seq level.

The *tme_cluster* function in IOBR is developed for profiling the tumor microenvironment, providing a clear depiction of a patient’s immune landscape. Utilizing data on TME cell infiltration and gene signature scores, *tme_cluster* performs unsupervised clustering to identify optimal TME patterns. To delve deeper into how intracellular signaling and intercellular interactions affect TME characteristics, we have developed the *lr_cal* function, based on the compute_LR_pairs from the easier package. This function calculates interaction scores for 813 LR pairs, using either count or TPM data. It aids in synthesizing a comprehensive overview of the TME by integrating various types of prior knowledge.

### Evaluating Mutations Associated With Specific Signatures and Delineating Pertinent Mutation Landscapes

Examining the TME, charting genomic alterations, and understanding their interrelated mechanisms are crucial for gaining insights that enhance patient classification and inform the development of personalized treatment strategies. It is critical to recognize that specific gene mutations can trigger the onset of cancer, affecting how cancer cells interact with the immune system and respond to immunotherapy across diverse cancer types. Addressing this aspect, IOBR has developed specialized functions for genomic analysis, catering to the intricate needs of understanding and interpreting these dynamics.

Within the IOBR framework, the *make_mut_matrix* function is designed to process genomic data in MAF and transform it into a format that is suitable for the *find_mutations* function. Utilizing *find_mutations* involve the simultaneous processing of genomic MAF data and a selected gene signature matrix. This function enables users to delve into the relationships between specific genes or signatures and mutations. We are thus equipped to investigate the differences in gene expression between wild-type and mutant samples of interest. Furthermore, IOBR allows for an intuitive examination of genomic alterations in samples with high and low gene or signature scores through a comprehensive tumor atlas, providing a clear visualization of the genomic landscape associated with these varying scores, such as oncoplot and boxplot.

### TME data visualization and Statistical analysis

Furthermore, to streamline the analysis process, IOBR incorporates the iobr_cor_plot function, designed for the efficient and rapid exploration of various datasets. This function dynamically generates statistical results and effectively illustrates the correlation between signatures and specific phenotypes, such as therapeutic responses or carcinogenic infection statuses. Additionally, IOBR offers the get_cor_matrix and get_cor functions, capable of describing the Pearson correlation between two or more features. The sig_forest function facilitates users to integrate the survival analysis output originated from the batch_surv function, and depicts a forest plot with hazard ratios of multiple signatures. Currently, leveraging signatures to predict specific phenotypes and survival benefits in response to therapy is a well-established approach in preclinical bioinformatics analysis. The sig_roc function, built on the pROC R package, is adept at outlining AUC curves for multiple signatures. The parameter compare method within this function enables users to assess the statistical difference between any two signatures of interest with an optional method. Additionally, the time_roc function, based on the timeROC R package, excels in creating time-independent ROC curves to evaluate the predictive performance of various variables.

The *sig_box* function can be employed to examine the correlation between a categorical variable and a specific signature, presenting a boxplot to show the statistical variance in signature scores across different categories. Meanwhile, the *sig_heatmap* function visually demonstrates the differences in correlations between features and categories through heatmaps.

Furthermore, IOBR efficiently visualizes relationships between signature genes and targeted variables (binary or continuous) using similar methods. It is also capable of identifying signatures significantly correlated with a specific signature of interest.

Considering that multiple signatures and signature genes may be enriched, IOBR includes a range of functions for batch statistical analysis and visualization. This encompasses batch survival analysis for either continuous signature scores or categorized phenotype subgroups, and the aforementioned batch correlation analysis employs statistical tests such as the Wilcoxon test and Partial Correlation Coefficient (PCC).

## Discussion

The rapid development and widespread application of sequencing technologies have enabled scientists to dissect the tumor’s immune microenvironment from multiple dimensions. Currently, RNA-seq has evolved into a mature and cost-effective analytical technique. It is increasingly utilized to explore cancer-immune interactions and characterize cancer cells and the tumor microenvironment. Computational methods using transcriptomic profiling are instrumental in understanding tumor immunity and in characterizing prognostic and predictive markers of immune therapy response [8].These methods provide valuable insights into immune response predictive markers, such as estimates of tumor-immune cell infiltration and gene expression signatures [29]. Furthermore, the advancement of single-cell sequencing technology offers high-resolution data on immune cell populations and the ability to detect variations between individual cells and cell groups [7, 30]. By integrating and analyzing multi-omic data, including transcriptomics, single-cell, and genomics, we can dissect the underlying gene regulatory mechanisms and unveil the landscape of the tumor microenvironment. This deepens our understanding of the interactions and biological mechanisms between immune and tumor cells [31, 32]. However, the complexity and growing volume of multi-omics data introduce new opportunities and challenges in analyzing the tumor microenvironment.

In previous research, our team developed IOBR, a user-friendly and comprehensive analysis tool oriented towards TME analysis [6]. IOBR assists users in efficiently and accurately parsing and visualizing the TME, exploring clinically relevant features and biomarkers. In this study, we have further expanded IOBR’s analysis and visualization capabilities. Building on a multi-omics approach with a focus on transcriptomics, IOBR comprehensively dissects the tumor immune microenvironment to unearth features related to tumor immunity and immune therapy responses. IOBR2 offers six main functional modules. It goes beyond effective systemic analysis of transcriptomic, genomic, and single-cell data by compiling a wide range of statistical and clinical analysis methods. Additionally, IOBR2 integrates an array of visualization functions, enhancing model construction and validation capabilities. IOBR2 provides a streamlined and efficient transcriptomic data preprocessing workflow, including data quality control and processing. An estimation function for signatures has been developed in IOBR2, alongside the integration of various cell deconvolution algorithms for rapid parsing and characterization of the TME. Besides the signatures documented in IOBR, IOBR2 has incorporated published single-cell signatures. It allows users to customize signatures and gene sets based on their findings in bulk RNA-seq or scRNA-seq data and their oncological insights. Users can then validate their findings in different transcriptomic datasets using IOBR2, including feature score computation and cell deconvolution. Furthermore, unveiling TME patterns and intra-tumor interactions, providing comprehensive descriptions of the TME, identifying effective biomarkers, and decoding the mechanisms behind patient treatment responses to ultimately predict immunotherapy efficacy have always been critical research directions [33]. Addressing this, IOBR2 has added a new module for TME interaction analysis. This module employs clustering to determine TME patterns and analyzes the interactions of infiltrating cell receptor-ligand pairs within the microenvironment, offering diverse perspectives on the TME. Additionally, IOBR2 provides various visualization functions suitable for different scenarios, facilitating the batch visualization of TME features and enabling rapid analysis of correlations between TME characteristics and clinical phenotypes. We have also introduced feature screening and model construction functions, assisting clinicians in swiftly identifying targets and biomarkers closely associated with patient treatment prognosis.

## Conclusion

Immunotherapy has revolutionized cancer treatment in recent years. The role of TME in these therapies is gaining increasing acknowledgment. The IOBR package offers a comprehensive downstream transcriptomic analysis process for tumor microenvironment analysis. It synergistically combines scRNA-seq and genomic data to present a multi-dimensional landscape of the TME. IOBR has multiple functions for microenvironment analysis, including cell abundance analysis and signature score calculation. It can also analyze TME interactions and integrate traditional analytical and modeling approaches, providing a comprehensive analysis and visualization solution for transcriptome projects. IOBR is expected to continue playing a significant role in the future with the ongoing advancements in multi-omics and artificial intelligence. It is ready to advance research in cancer immunology and immuno-oncology, providing new understanding of tumor immunity and responses to immunotherapy.

## Acknowledgments

The work is supported by Guangdong Provincial Science and Technology Project (No. 2020A0505090007 to J. Wang). The authors thank EGA and David for providing the multi-omics data of IMvigor210. The authors thank all the patients, their families, investigators, and healthcare professionals who participated in this study. The authors are also grateful to the researchers who generated and shared the sequencing data openly.

## Authors’ contributions

DQZ contributed to the conceptualization and study design. DQZ, YRF, PL and WJQ contributed to acquisition. DQZ and QQM contributed to data analysis and interpretation. DQZ, YRF, WJQ, QQM and SXW contributed to package development. DQZ, and YRF drafted the tutorial. DQZ, YRF and XX were responsible for drafting the initial draft of the manuscript. DQZ, YRF, PL, GCY and SXH participated in revising the manuscript. WJL and GCY supervised the project and reviewed the manuscript. All the authors have read, discussed, and approved the final version of the manuscript. The corresponding author had full access to the data in the study and took responsibility for the integrity of the data and the accuracy of the data analysis.

## Ethics approval and consent to participate

Ethics approval and patient-informed consent for The Cancer Genome Atlas (TCGA), Gene Expression Omnibus (GEO), and European Genome-phenome Archive (EGA) were waived due to their public availability.

## Data and code availability

The multi-omics data retrieved from a trial of atezolizumab for bladder cancer (IMvigor210) were available in the European Genome-Phenome Archive database under accession codes EGAS00001002556 [2]. Two normalized microarray data for gastric cancer datasets GSE62254 and GSE57303 were downloaded from the GEO database (https://www.ncbi.nlm.nih.gov/geo/). RNA-seq data for TCGA-STAD were obtained from The Cancer Genome Atlas (TCGA, https://portal.gdc.cancer.gov/) via the University of California, Santa Cruz Xena platform (UCSC Xena, https://gdc.xenahubs.net/).

The IOBR R package is available at https://github.com/IOBR/IOBR [6]. The gitbook (https://iobr.github.io/book/) provides a complete analysis workflow for each module within the package, including numerous examples and detailed explanations of its functions. Any additional information required to reanalyze the data reported in this paper is available from the lead contact upon request.

## Consent for publication

Not applicable.

